# Arterial iron regulates vasodilation during anemia via endothelial holo-alpha globin

**DOI:** 10.64898/2025.12.09.693297

**Authors:** Luke S. Dunaway, Shruthi Nyshadham, Wyatt J. Schug, Skylar Loeb, Brooke O’Donnel, Zuzanna Juskiewicz, Nasim Abib, Melissa A. Luse, Macy Stahl, Timothy M. Sveeggen, Pooneh Bagher, Jason Allen, Adam N. Goldfarb, Brant E. Isakson

## Abstract

Iron deficiency is a highly prevalent nutrient deficiency and the most common cause of anemia. Although iron deficiency exacerbates cardiovascular disease, the direct impact of iron deficiency on the vasculature remains unstudied. We assessed iron levels across the vascular endothelium and found resistance artery endothelial cells to have the lowest iron stores suggesting they may be especially impacted by iron deficiency. Anemia has previously been shown to increase arterial NO signaling in patients, and we have previously shown endothelial α-globin (Hbα) scavenges nitric oxide (NO) in the resistance artery endothelium. We hypothesize iron regulates vascular function through downregulation of endothelial Hbα. To test this, we used a novel model of iron deficiency anemia (IDA). In female mice, IDA increased NO signaling which was rescued to control levels by repletion of vascular iron with ferric dextran. Despite being similarly anemic and having a similar reduction in Hbα protein, there were no changes in NO signaling across groups in male mice. We further measured whether Hbα was in its heme-bound (holo-Hbα) or heme-free (apo-Hbα) state and found males did not fully lose holo-Hbα. Using endothelial specific Hbα knockout mice, we show loss of endothelial Hbα is necessary for increased NO signaling in IDA and for the rescue of NO signaling by ferric dextran in female mice. Altogether the data presented here demonstrate iron modulates endothelial NO signaling through the regulation of Hbα.

## Introduction

Iron deficiency is a common nutrient deficiency and is associated with chronic diseases such as heart failure^1^, chronic kidney disease^2^, and other diseases with chronic inflammation^3^. Each of these diseases is also associated with endothelial dysfunction. The endothelium regulates diverse processes such as vascular resistance, tissue perfusion, immune cell extravasation, and waste removal which requires specialization of endothelial phenotypes along the vascular tree. Iron is an important cofactor for proteins involved in transcriptional regulation, metabolism, and redox signaling^4^. As such, iron likely plays an important role in endothelial biology, but the role of iron on endothelial phenotypes remains unstudied.

Perhaps the most central endothelial function is the regulation of tissue perfusion. Endothelial cells (ECs) modulate hemodynamics to ensure adequate nutrient and oxygen delivery to tissues. Previous work investigating hemodynamic consequences in iron deficiency has focused on the impact of anemia and left the direct consequences of vascular iron deficiency unexplored. Anemia has been previously shown to increase nitric oxide (NO) signaling^5^. This has been attributed to changes in blood hemoglobin as hemoblobin is a potent NO scavenger. However, ex vivo and in vitro studies demonstrate hemoglobin in intact red blood cells is a ∼1000-fold worse scavenger of NO compared to free hemoglobin and has minimal impact on arterial NO signaling in perfused preparations^6^. This suggests changes in hematocrit in anemia may be insufficient to increase NO signaling. The alpha chain of hemoglobin (α-globin; Hbα) is also expressed in the endothelium of resistance arteries, the arteries which control blood flow^7–10^. Here Hbα binds directly to endothelial NO synthase and scavenges the NO it produces. Because hemoglobin is tightly regulated by systemic iron homeostasis in erythrocytes, we hypothesize vascular iron deficiency downregulates Hbα thereby increasing NO signaling in iron deficiency anemia (IDA).

## Methods

### Single cell RNA Sequencing

Previously published single cell RNA sequencing data of isolated mouse ECs^11^ (GSE235192) and human ECs^12^ (https://www.vascularcellatlas.org/) were reanalyzed in Rstudio (4.1.2) with Seurat (4.3.0). For mouse ECs, cells from normal chow fed mice were subset for analysis. For human ECs, organs without representation of all four endothelial subtypes were excluded. The final dataset included adipose, bladder, heart, large intestine, lung, muscle, esophagus, pancreas, small intestine, thymus, trachea. The categorization of arterial, capillary, venous, and lymphatic ECs were kept as in the original manuscripts. Iron dependent proteins^4^ that were expressed in >5% were considered for analysis.

### En Face

Mesenteric arteries: Third order mesenteric arteries were excised and cleaned of adipose and connective tissue. Arteries were cut to length and pinned out using .0005” tungsten wire (ElectronTubeStore) onto polymerized Sylgard 184 (Electron Microscopy Sciences). Fixation was performed using 50:50 acetone/methanol for 8 minutes on ice, followed by three 5-minute washes in 1X PBS. Arteries were then cut *en face* using microdissection scissors and pinned flat with the lumen facing upward using tungsten wire.

Thoracic aortas: Thoracic aortas were isolated and cleaned of adipose and connective tissue. Fixation was carried out in two steps: arteries were first incubated in 4% paraformaldehyde (PFA) for 30 minutes on ice, followed by incubation in 0.4% PFA overnight at 4°C. Samples were then washed three times in 1X PBS for 5 minutes each. Aortas were kept intact during permeabilization and immunostaining and were cut *en face* using microdissection scissors on the microscope slide immediately before mounting.

Permeabilization and immunostaining: Samples were permeabilized in 0.2% NP-40 in PBS for 30 minutes at room temperature and then blocked in 5% bovine serum albumin (BSA) in 0.2% NP-40/PBS for 1 hour. Primary antibody staining was performed in 0.5% BSA in 0.2% NP-40/PBS using the following antibodies: ferritin light chain (ab69090, 1:100, Abcam) and claudin 5 (35-2500, 1:100, Invitrogen). After three 5-minute washes in 1X PBS, samples were incubated with secondary antibodies (1:400 in 0.5% BSA in 0.2% NP-40/PBS) for 1 hour at room temperature. Nuclei were stained with DAPI (Invitrogen; D1306; final concentration, 0.1 mg/mL). Samples were mounted on glass slides using Prolong Gold Antifade Mountant (Invitrogen; P36930). Images were acquired using an Olympus FV3000 confocal microscope equipped with a 60X oil immersion objective. Postprocessing was performed using FIJI.

### Cell culture models

To deplete iron, human coronary artery ECs and human dermal lymphatic ECs were grown to confluence and treated with 100 μM deferoxamine (DFO; MedChemExpress) for 24 hours. RNA was isolated with Quick-RNA MiniPrep (Zymo Research). RNA sequencing and transcriptomic analysis was performed as described below.

To model resistance arteries, endothelial cells were cultured following the vascular cell co-culture model as previously reported^13^. Briefly, human coronary artery smooth muscle cells (SMCs) were plated (75,000 cells/well) on the bottom of a 6-well plate or on the underside of the transwell. The following day, human coronary artery ECs were seeded on the top side of the transwell. An additional group of ECs were plated in a transwell dish without smooth muscle cells. In all three conditions, endothelial cell growth media MV (Promocell) was placed on the top of the transwell and SMC media (Lonza) was placed in the bottom chamber of the transwell. Two days after plating ECs, the media was changed, and ECs were collected the following day. For transcriptomics, RNA was isolated using Quick-RNA MiniPrep (Zymo Research). Protein was collected as described for western blot below.

To assess the labile iron pool, ECs were stained with FerroOrange (Cayman Chemical). ECs were rinsed 3 times in HBSS and incubated in HBSS with 1 μM FerroOrange for 30 min at 37C 5% CO_2_. Cells were dissociated with trypsin, centrifuged 1000 x g for 3 min, and suspended in 1 μM FerroOrange, 1 μM sytox blue in HBSS. Samples were run using a Cytek Aurora 5L Spectral Flow Cytometer. Cells positive for Sytox Blue were excluded and the mean fluorescent intensity (MFI) of FerroOrange was calculated using FCS Express, Version 7 (De Novo Software).

### Bulk RNA Sequencing

RNA was isolated from cultured ECs. PE150 reads were generated with the Illumina NovoSeq platform. The paired-end reads were mapped to the hg38 reference genome using STAR (v2.7.9a). Gene counts were generated with featureCounts in the Subread package. For previously published data of human umbilical vein endothelial cells exposed to shear stress, raw counts were downloaded from the NCBI Gene Expression Omnibus (GEO; GSE158081). For all datasets, DESeq2 (v1.34.0) was used to identify differentially expressed genes (DEGs) (Padj < 0.05). Ensemble IDs were converted to gene symbols using the org.Hs.eg.db package.

### Model of Iron Deficiency Anemia

All experiments were approved by the University of Virginia Animal Care and Use Committee and followed the National Institutes of Health guidelines for the care and use of laboratory animals. Endothelial *Hba1* knockout (*Cdh5-CreERT2^+/-^ Hba1^fl/fl^*) mice were generated as previously reported^9^. Cre^+^ and Cre^-^ littermates received tamoxifen injections (100 μL at 10 μg/mL in peanut oil) were administered for 10 days starting at 6 weeks of age. To model IDA, male and female mice were fed an iron deficient (2-6 ppm; TD.80396) beginning at 3 weeks of age. Phlebotomies of 10% blood volume were performed at 9 and 11 weeks to drive the progression of anemia. A subset of mice (IDA+FeDex) received an injection of iron dextran (20 mg/kg; Sigma). Control mice were fed a nutrient matched control (48ppm; TD.220330) and received an injection of iron dextran at 12 weeks. All experiments were performed 7-8 days after injection. Body composition was measured by EchoMRI Body Composition Analyzer (EchoMRI^TM^). Terminal blood collections were taken from the heart. Complete blood counts were analyzed on a VETSCAN HM5 hematology analyzer (Abaxis).

### Laser speckle contrast imaging

To assess NO signaling, blood flow was assessed with PeriCam PSI (PeriMed) similar to previously published methods^14,15^. The mice were anaesthetized with 4% isoflurane delivered with room air then reduced to 1.5%. Core body temperature was maintained between 37.0-37.4°C. After at least 15 min equilibration, blood flow was measured in both hind paws of the mice before and after injection with nitro-l-arginine methyl ester hydrochloride (L-NAME; 100 mg/kg). The response to L-NAME has been used as a measure of NO signaling in vivo in mice and humans^5,16^.

### Liver iron assay

Iron assay kit (Sigma MAK025) was used to assess liver iron. Liver samples were taken from mice perfused with PBS and dehydrated in a 65 °C incubator for 48 hours. The livers were crushed with a mortar and pestle and weighed. Livers were sonicated in iron assay buffer and centrifuged at 16,000 x g for 10 min at 4 °C. Iron was assessed following the manufacturer protocol and normalized to dry weight of the sample.

### Western blots

Cells and tissues were lysed in RIPA buffer (50mmol/L Tris-HCL, 150mmol/L NaCl, 5mmol/L EDTA,1% deoxycholate, 1% Triton-X100 in PBS pH 7.4) with protease (Sigma P8350) and phosphatase inhibitor cocktails (Sigma P5726 and P0044). Cells were sonicated 10x on setting 3 on Sonic Dimembrator Model 100 (Fisher Scientific) and centrifuged at 1000 x g for 5 min at 4°C. Isolated mesenteric arteries were crushed in liquid nitrogen and sonicated setting 3 and centrifuged at 1000 x g for 5 min at 4 °C. Protein concentration was determined by Pierce BCA Protein Assay (Thermo Fisher). Lysates were electrophoresed on 8% or 4-12-% Bis-Tris gels and transferred to nitrocellulose membranes. Loading was assessed by REVERT total protein stain following the manufacturer protocol. Membranes were blocked with 3% BSA at room temperature for 1 hour and stained with antibodies against FTH (Cell Signaling 4393S), FTL (abcam ab69090), TfR1(Thermo Fisher 13-6800), eNOS (BD Transduction 610297), eNOS p-S1177 (Cell Signaling 9571), Hbα (Thermo Fisher PA5-97820) followed by incubation with goat-anti-mouse or goat-anti-rabbit secondary antibodies (LI-COR Biosciences). Membranes were imaged on Odyssey CLx Infrared Imaging System (LI-COR Biosciences) and densitometry was quantified with LI-COR Image Studio.

### Holo-hbα assay

To have sufficient protein for the assay, renal vessels were isolated as previously described^16^. Briefly, perfused kidneys were place in ice cold PBS between two 100 μm filters and gently grated to remove the epithelium. Tissue was homogenized in immunoprecipitation buffer (20 mM Tris base, 130 mM NaCl, 0.5% Trion X-100, 1 mM EDTA, 1 mM EGTA, pH 7.5) supplemented with protease (Sigma P8350) and phosphatase inhibitor cocktails (Sigma P5726 and P0044). Homogenization was performed with a FastPrep-24 (MP Biomedicals) and sonicated on setting 3. Lysates were incubated at 4 °C for 2 hours rocking and centrifuged 1000 x g for 5 min at 4 °C. Protein concentration was determined by Peirce BCA assay (Thermo Fisher). 100 ug of protein was incubated with 4 μg anti-Hbα (Thermo Fisher 14537-1-AP) overnight at 4 °C rocking. Sheep anti-rabbit Dynabeads (Thermo Fisher) beads were washed 3X in blocking buffer (0.5% BSA, 0.2% fish skin gelatin, in PBS, pH 7.4) for 5 min at 4 °C before incubating with the lysate and antibody solution for 2 hours at 4 °C while rocking. After incubation, the supernatant was removed and beads were washed 2X with ice cold 0.05% Tween-20 in tris buffered saline. To elute the immunoprecipitated Hbα, beads were incubated in 0.1 M glycine (pH 2.3) for 15 min at room temperature rocking. A portion of the eluate was taken for immunoblotting and neutralized with 1 M Tris base (pH = 8.5) to a final concentration of 100 mM Tris base. Western blot was performed as described above. The remainder of the sample was used to assess the amount of heme pulled down.

To measure heme, samples and standards were divided into two tubes and mixed 1:1 with 2 mM oxalic acid. One set were incubated at 100 °C for 30 min to convert the heme to protoporphyrin IX. The other was kept at room temperature for 30 min. Protoporphyrin IX fluorescence was assessed in a plate reader (ex: 400 nm, em 620 nm). Heme concentration was calculated by subtracting out the respective unboiled sample then using the equation generate by the standard curve. Each sample was normalized to the amount of immunoprecipitated Hbα as assessed by western blot.

### Statistical analysis

Data are represented as means ± SEM. Statistical differences were analyzed by Student’s t-test, one-way or 2-way ANOVA followed by Holm-Sidak post hoc test where indicated. P < 0.05 were considered significant. Statistical analysis was performed using GraphPad Prism Software (version 10.1.0).

## Results

### Resistance artery endothelial cells are iron deplete

To better understand how iron deficiency impacts endothelial biology, we investigated where iron was localized across the microvasculature. Using our previously published single cell RNA sequencing (scRNA-seq) data set of ECs isolated from the mesentery and adipose tissue we analyzed the expression of *Tfrc* as a measure of labile iron stores. Using human coronary artery ECs, we confirmed *TFRC* is inversely correlated to iron in the endothelium (**Supplemental Figure 1A**). Regardless of the tissue, we found *Tfrc* was highest in the arterial endothelium and is lowest in the lymphatic endothelium (**Figure 1A**). We also investigated ferritin expression as a measure of iron stores. We found ferritin was lowest in the arterial endothelium and is highest in the lymphatic endothelium (**Figure 1B**). To investigate this in the human endothelium, we utilized a published scRNA-seq dataset of human ECs pooled from 11 different organs^12^. Similar to what we observed in mice, ferritin expression is low in arterial ECs and is highest in the lymphatic (**Figure 1C**) Notably, the differential regulation of iron across endothelial subtypes does not correlate with the expression of iron dependent proteins in either tissue (**Figure 1D**).

**Figure 1.**
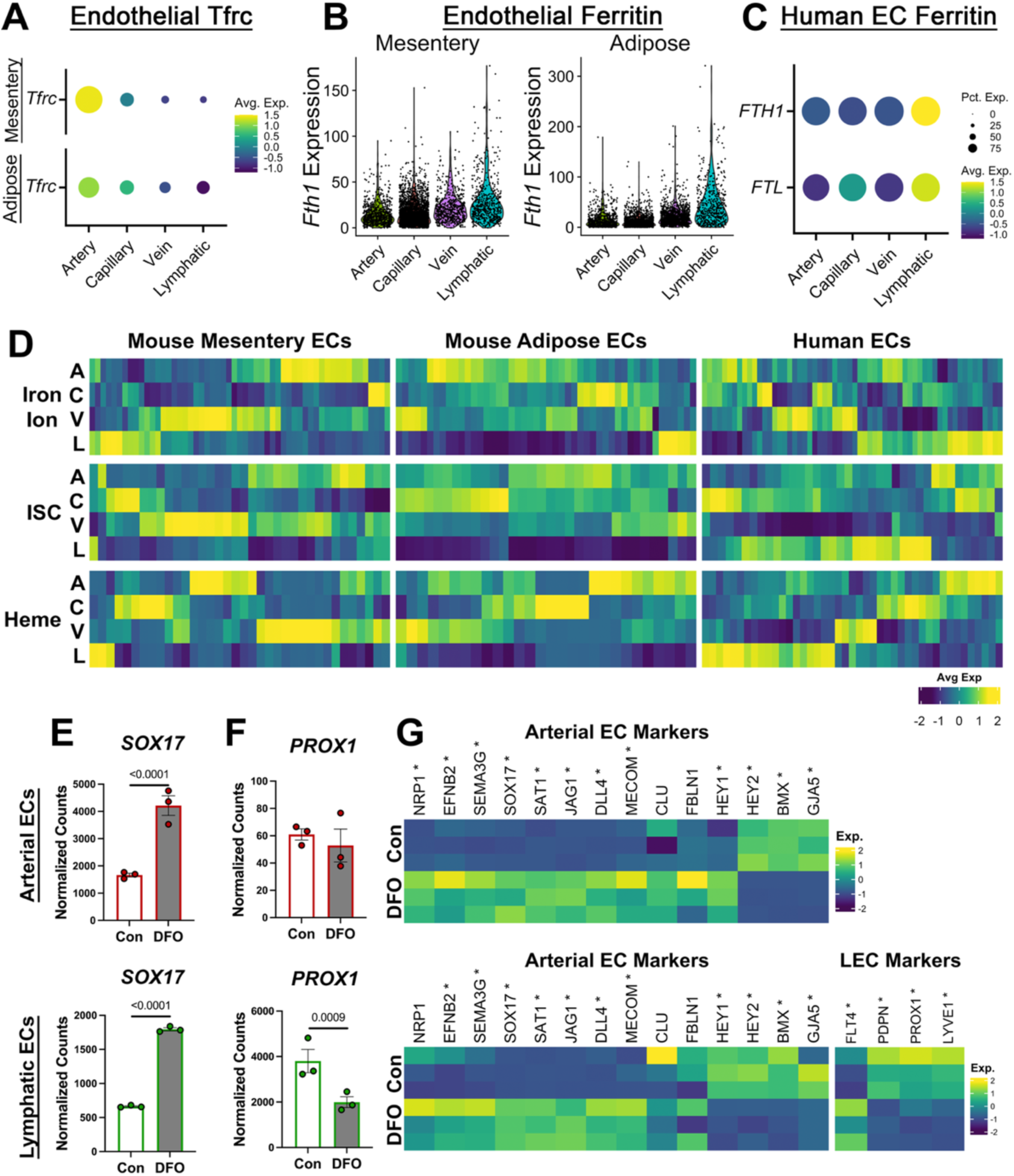
In the microvasculature, the arterial endothelium is iron deplete. (A-B) Transferrin receptor (*Tfrc*) and ferritin heavy chain (*Fth1;Fth1*) expression in previously published single cell RNA sequencing (scRNA-seq) data of endothelial cells (ECs) isolated from adipose and mesentery. (C) Expression of FTH1 and FTL in previously published scRNA-seq data of human ECs. (D) The expression genes encoding proteins dependent upon iron ions, iron sulfur clusters (ISC), and heme across endothelial subtypes. (E-G) Human coronary artery ECs and human dermal lymphatic ECs were treated with deferoxamine (DFO) for 2h hours. The expression of SOX17 (E) PROX1 (F) and markers of arterial and lymphatic EC phenotypes (G) were assessed by bulk RNA-seq. Statistical analysis was performed with DESeq2.

We next investigated if the low iron in the arterial endothelium is related to endothelial identity. To test this, we treated human artery ECs and human lymphatic ECs with deferoxamine (DFO) to chelate iron and assessed phenotypic changes by bulk RNA seq (**Figure 1E-G; Supplemental Tables 1-3**). *SOX17*, a transcription factor which promotes arterial EC identity, was upregulated in both cell types (**Figure 1E**). *PROX1*, a transcription factor which promotes lymphatic EC identity, is minimally expressed in arterial ECs and was downregulated in lymphatic ECs (**Figure 1F**). Assessing the expression of genes which have been previously used to assess endothelial identity^17^, we found a majority of arterial markers to be upregulated by iron chelation in both cell types and lymphatic EC makers to be downregulated in lymphatic ECs (**Figure 1G**). These data demonstrate an importance for low endothelial iron in arterial EC identity in the microvasculature.

Microvascular artery ECs have distinct biology compared to large conduit arteries^18^. For this reason, we compared FTL expression in conduit vs resistance arteries prepared *en face* to examine if iron stores are uniformly low across the arterial endothelium. Using aorta as a representative conduit artery and third order mesenteric arteries as a representative resistance artery, we found ferritin expression to be higher in the conduit artery endothelium compared to the resistance artery endothelium (**Figure 2A**). Biological differences between conduit and resistance arterial endothelium can be due to either the presence of myoendothelial junctions (MEJs) in resistance arteries, and/or the high flow rates in conduit arteries (**Figure 2B**). To investigate the contribution of MEJs, we used a vascular cell co-culture model previously established by our lab which promotes the formation of MEJs and recapitulates the cell biology of resistance artery ECs^7,13,19^. We cultured human coronary artery ECs in three conditions: cultured without SMCs (monolayer), co-cultured without direct contact with SMCs (no-contact), and co-cultured with SMCs such that MEJs could form (contact) (**Figure 2C**). Bulk RNA seq of ECs grown in these three conditions found over 2700 genes were differentially regulated demonstrating communication with SMCs has broad implications for EC phenotypes. (**Figure 2D; Supplemental Tables 4-7**). Labile iron was directly assessed by FerroOrange and we found contact with SMCs does not impact endothelial iron (**Figure 2E**). We also found no evidence of differential iron regulation as assessed by western blot of transferrin receptor, ferritin heavy chain, and ferritin light chain. (**Figure 2F**). To check if sheer was responsible for the difference in iron regulation across the arterial endothelium, we reanalyized published bulk RNA-seq data of ECs exposed to continuous shear stress (11.5 dynes/cm^2^) for up to 48 h revealed >4300 differentially expressed genes (**Figure 2G-H; Supplemental Tables 9-12**)^20^. In these experiments, we found shear stress increased *FTH1* and *FTL* and decreased *TFRC* (**Figure 2I**) and thus is likely responsible for differences across the arterial endothelium. Altogether our findings demonstrate the resistance artery endothelium is the most basally iron deplete across the vasculature, suggesting it may be especially susceptible to changes in iron homeostasis (**Figure 2J**).

**Figure 2.**
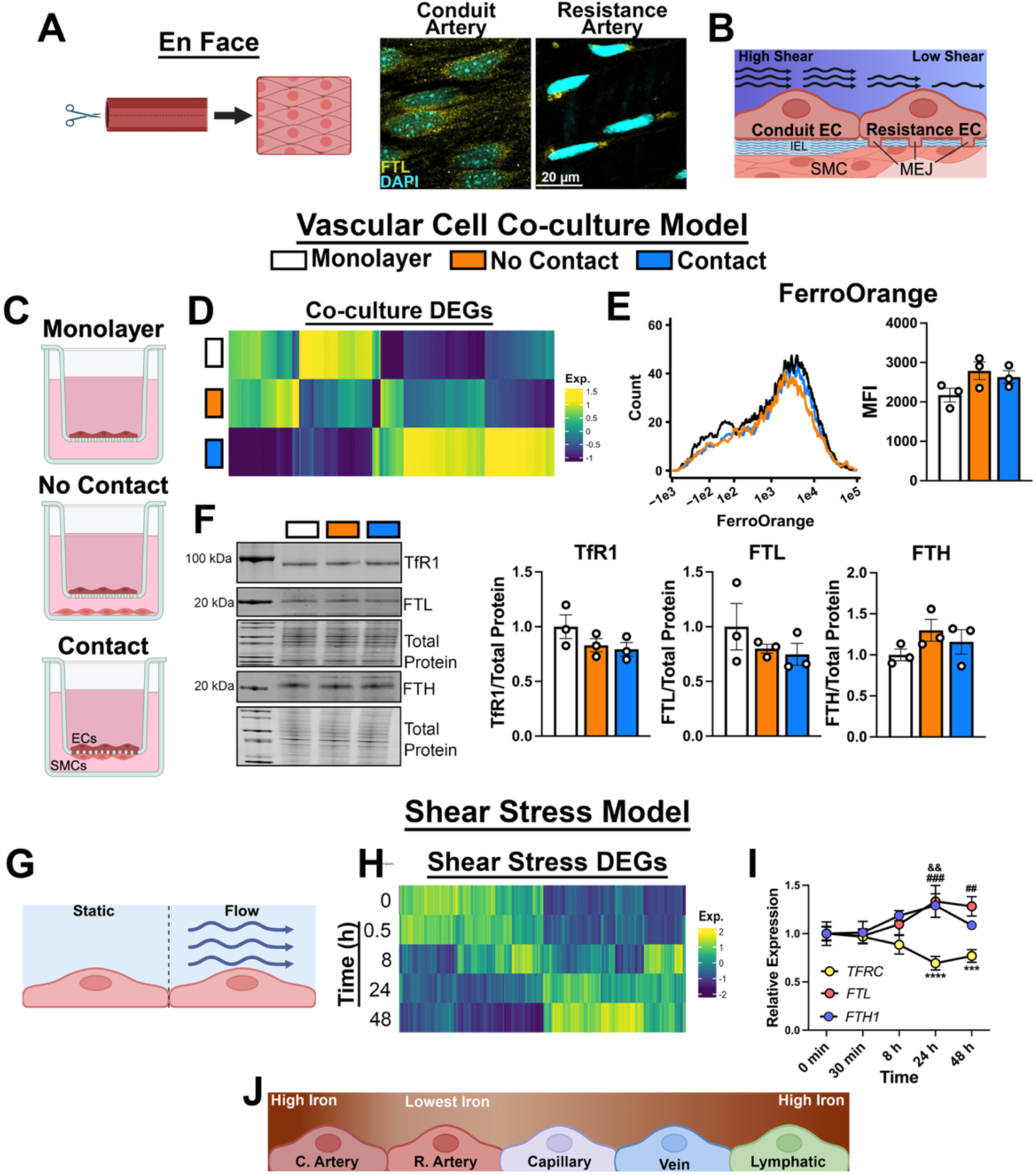
Shear stress dictates intracellular iron in conduit and resistance arteries. (A) *En face* imaging ferritin light chain (FTL) in a conduit artery (aorta) and resistance artery (3° mesenteric artery). (B) Schematic highlighting shear stress and myoendothelial junctions (MEJ) as key determinants of conduit and resistance artery endothelial biology. (C-F) Human coronary artery endothelial cells (ECs) were grown in transwells either in a monolayer, co-cultured with smooth muscle cells (SMCs) without direct contact (no contact), or co-cultured in direct contact with SMCs (contact). (D) Heatmap of differentially expressed genes (DEGs) across all three conditions as assessed by bulk RNA sequencing. (E) Labile iron was assessed by flow cytometry with FerroOrange. (F) Transferrin receptor (*TFRC*; TfR1), ferritin light chain (*FTL*; FTL) and ferritin heavy chain (*FTH1*; FTH) were assessed by western blot. (G-I) Reanalysis of human umbilical vein ECs grown in static or flow conditions for up to 48 h (H) Heatmap of genes differentially regulated by shear stress. (I) *TFRC*, *FTL*, and *FTH1* expression in static and flow conditions. ***p<0.001 vs *TFRC* 0 min; ****p<0.0001 vs *TFRC* 0min; &&p<0.01 vs *FTH1* 0 min; ##p<0.01 vs *FTL* 0 min; ###p<0.001 vs *FTL* 0 min (J) Schematic summarizing iron across the vascular endothelium. IEL: internal elastic lamina; C. Artery: conduit artery; R. Artery: resistance artery.

### Iron deficiency anemia increases nitric oxide signaling

The resistance artery endothelium plays a central role in the regulation of blood flow through the production of vasoactive compounds such as NO, and it has been previously shown that patients with IDA have altered hemodynamics due to elevated NO signaling. To investigate if changes in endothelial iron contribute to the increased NO, we created a mouse model of IDA with replete vascular iron. Mice were placed on an iron deficient diet at weaning and phlebotomies were performed when the mice were 9 and 11 weeks old. Mice then received in intraperitoneal injection of saline or ferric dextran (FeDex; 20 mg/kg) to replete vascular iron (**Figure 3A**). Both male and female IDA mice had decreased liver iron demonstrating iron deficiency as well as blood hemoglobin, mean corpuscular hemoglobin concentration (MCHC), and hematocrit demonstrating anemia. (**Figure 3B-C**) Iron dextran did not rescue anemia in these mice. Arterial iron stores were assessed by FTL expression in isolated mesenteric arteries. IDA depleted vascular iron and was replete with ferric dextran. (**Figure 3D**) This results in a mouse model with anemia but replete for vascular iron.

**Figure 3.**
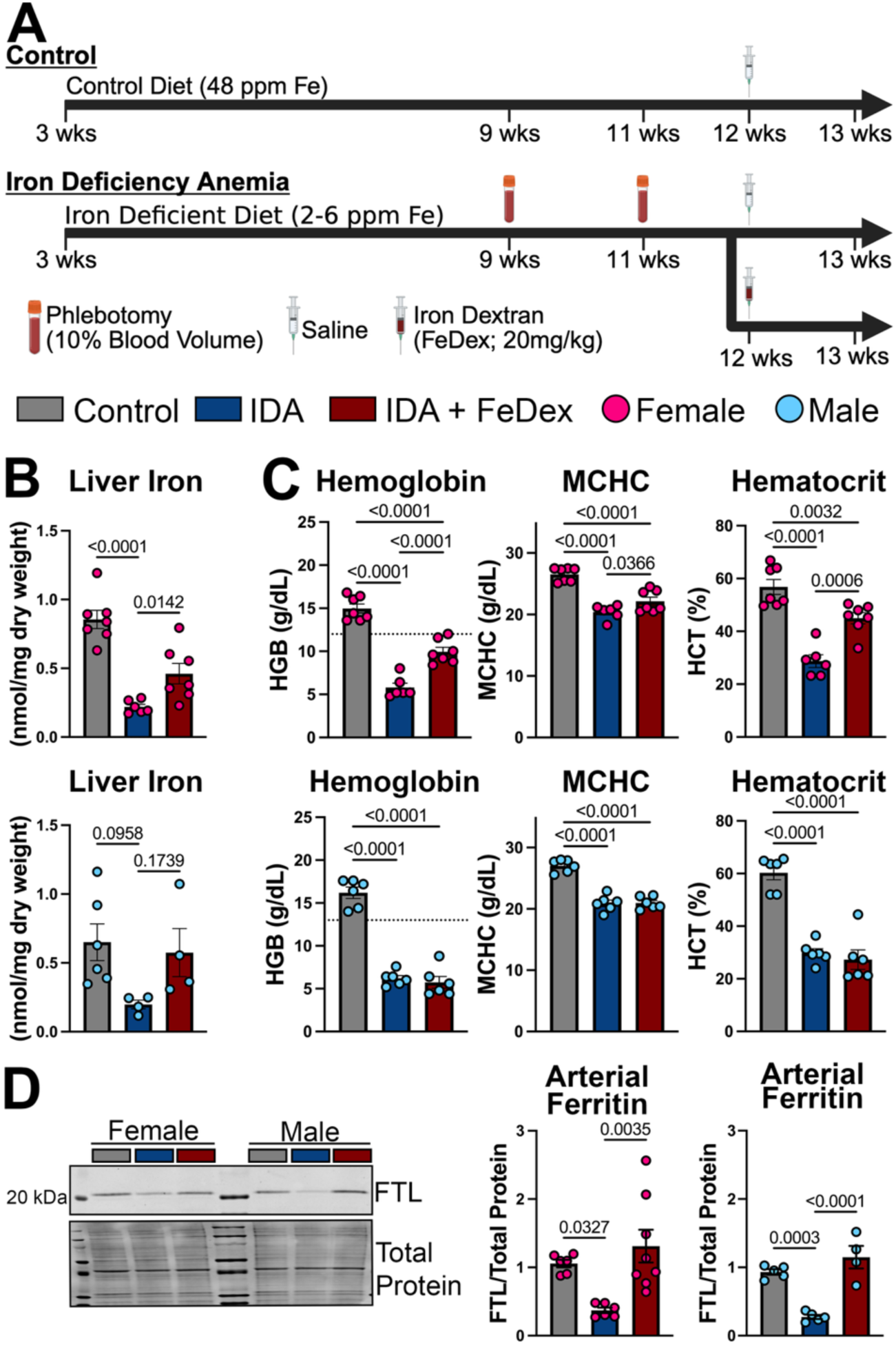
Model of iron deficiency anemia with replete vascular iron. (A-D) Female (pink) and male (blue) mice were fed control (48 ppm Fe) or iron deficient diet (2-6 ppm Fe) beginning at weaning. Phlebotomies were performed in iron deficient mice at 9 weeks and 11 weeks to aid in the progression of anemia then received a single injection of iron dextran (FeDex; 20 mg/kg) or vehicle (saline) at 11 weeks. All experiments were performed at 13 weeks. (B) Liver iron assessed by colorimetric assay. (C) Hemoglobin, mean corpuscular hemoglobin concentration (MCHC), and hematocrit from complete blood counts. Dashed line notes cut off for anemia. (D) Ferritin light chain (FTL) assessed by western blot in isolated mesenteric arteries. Statistical significance was determinized by on-way ANOVA and Holms-Sidac post hoc analysis. IDA: iron deficiency anemia; LCSI: laser speckle contrast imaging

We used laser speckle contrast imaging (LSCI) to assess the blood flow response to L-NAME as a measure of NO signaling in these mice. The response to a NOS inhibitor such as L-NAME is commonly used to assess NO signaling in mice and humans where a larger response indicates higher basal NO signaling^5,16^. We found female IDA mice to have higher NO signaling whereas mice with replete vascular iron had a similar response to L-NAME as control mice (**Figure 4A**). In male mice, we observed no differences in NO signaling between groups (**Figure 4B)**.

**Figure 4.**
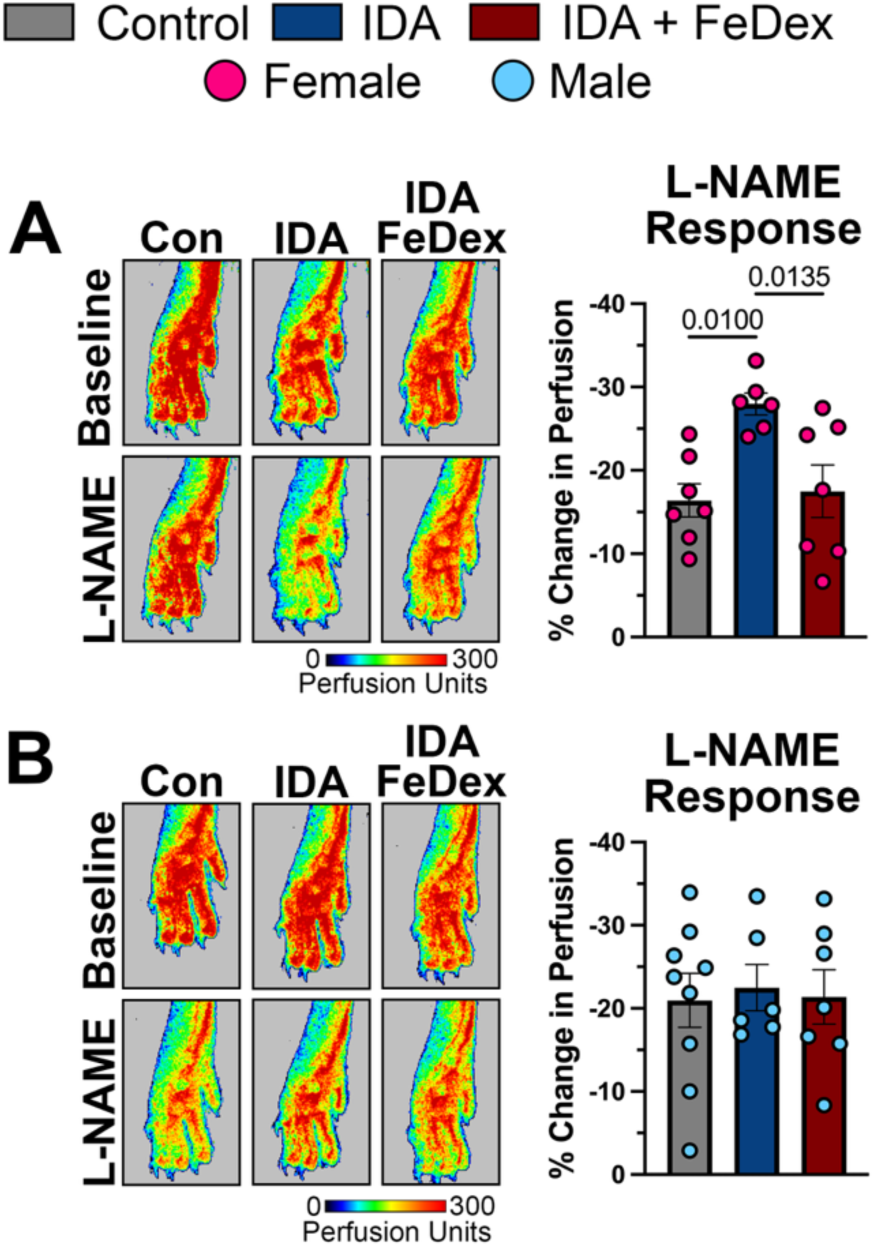
Vascular iron deficiency anemia increases endothelial nitric oxide. (A-B) Changes in blood flow in hind paws of female (A) and male (B) mice were assessed by laser speckle contrast imaging in response to intraperitoneal injection of the nitric oxide synthase inhibitor, L-NAME. Statistical significance was determinized by on-way ANOVA and Holms-Sidac post hoc analysis.

### Loss of endothelial holo-Hbα increases NO in IDA

NO signaling may be increased by either increased NO production or a decrease in NO scavenging. We assessed the NO metabolites, nitrate and nitrite, in the plasma and observed no change in plasma nitrate and nitrite across groups (**Figure 5A-B**). Similarly, we found IDA did not increase eNOS expression or phosphorylation at the activating site S1177 (**Figure 5C-D**). We have previously shown Hbα is expressed in the endothelium and inhibits NO signaling by scavenging NO. We measured Hbα expression in isolated arteries and found IDA decreased Hbα in female and male mice and FeDex rescued Hbα in female mice (**Figure 5E**). Heme proteins can either exist in their apo (heme-free) or holo (heme-bound) state. To assess holo-Hbα in our model, Hbα was immunoprecipitated out of isolated arteries and the amount of heme pulled down was measured. We pulled down equal amounts of Hbα in males and females. In females we found holo-hα was at the detection limit for the assay and increased when treated with FeDex. In males, we found a much larger proportion of holo-hα in controls which was decreased in IDA (**Figure 5F**). Together this suggests that while both males and females lose Hbα expression in IDA, females start with a very small pool of holo-hα that is easily lost in IDA and increased with FeDex. Males start with a large pool of holo-Hbα in the endothelium, which although decreased in IDA, is still very much detectable (**Figure 5G**).

**Figure 5.**
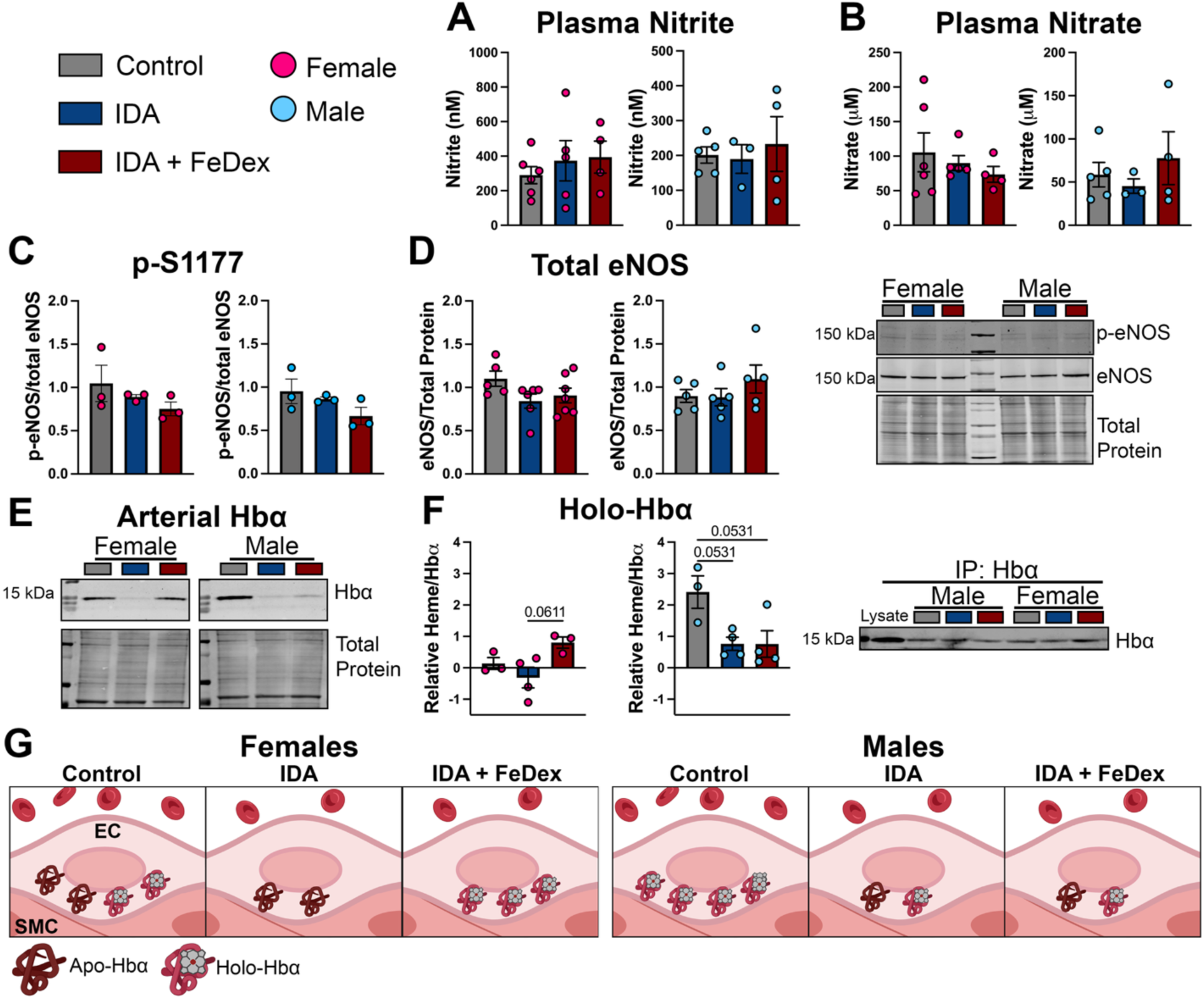
Vascular iron regulates endothelial a-globin, but not endothelial nitric oxide synthase. (A-B) Plasma nitrate and nitrite. (C-D) S1177 phosphorylation and total endothelial nitric oxide synthase (eNOS) assessed by western blot. (E) Endothelial α-globin (Hbα) expression assessed by western blot. (F) Hbα was immunoprecipitated and heme was measured in the immunoprecipitate to assess holo-Hbα. (G) Summary of Hbα and holo-Hbα levels in each condition. Statistical significance was determinized by on-way ANOVA and Holms-Sidac post hoc analysis. IDA: iron deficiency anemia; FeDex: iron dextran.

Based on these results, we hypothesized that the increased NO in females, is due to the loss of functional Hbα and that males do not increase NO in IDA because, although decreased, a functional pool of Hbα still remains. To test this hypothesis, we utilized endothelial specific Hbα knockout mice (*Cdh5-CreERT2^+/-^ Hba1^fl/fl^*) which we have previously shown selectively depletes Hbα from the endothelium. Deletion of *Hba1* in the endothelium does not impact blood hemoglobin levels or arterial iron stores (**Figure 6A-B**). We again measured NO signaling using LSCI. In female Cre^-^ mice, we found IDA increased NO signaling which was rescued by repleting vascular iron. However in Cre^+^ mice, NO signaling was not rescued by repletion of vascular iron demonstrating iron regulates arterial NO signaling through endothelial Hbα. Similar to what was observed in male wildtype mice, male Cre^-^ mice had no change in NO signaling in response to IDA. Whereas male Cre^+^ mice had increased NO signaling which was not rescued by FeDex phenocopying female Cre+ mice. These data demonstrate the loss of Hbα is necessary for increased NO signaling in IDA.

**Figure 6.**
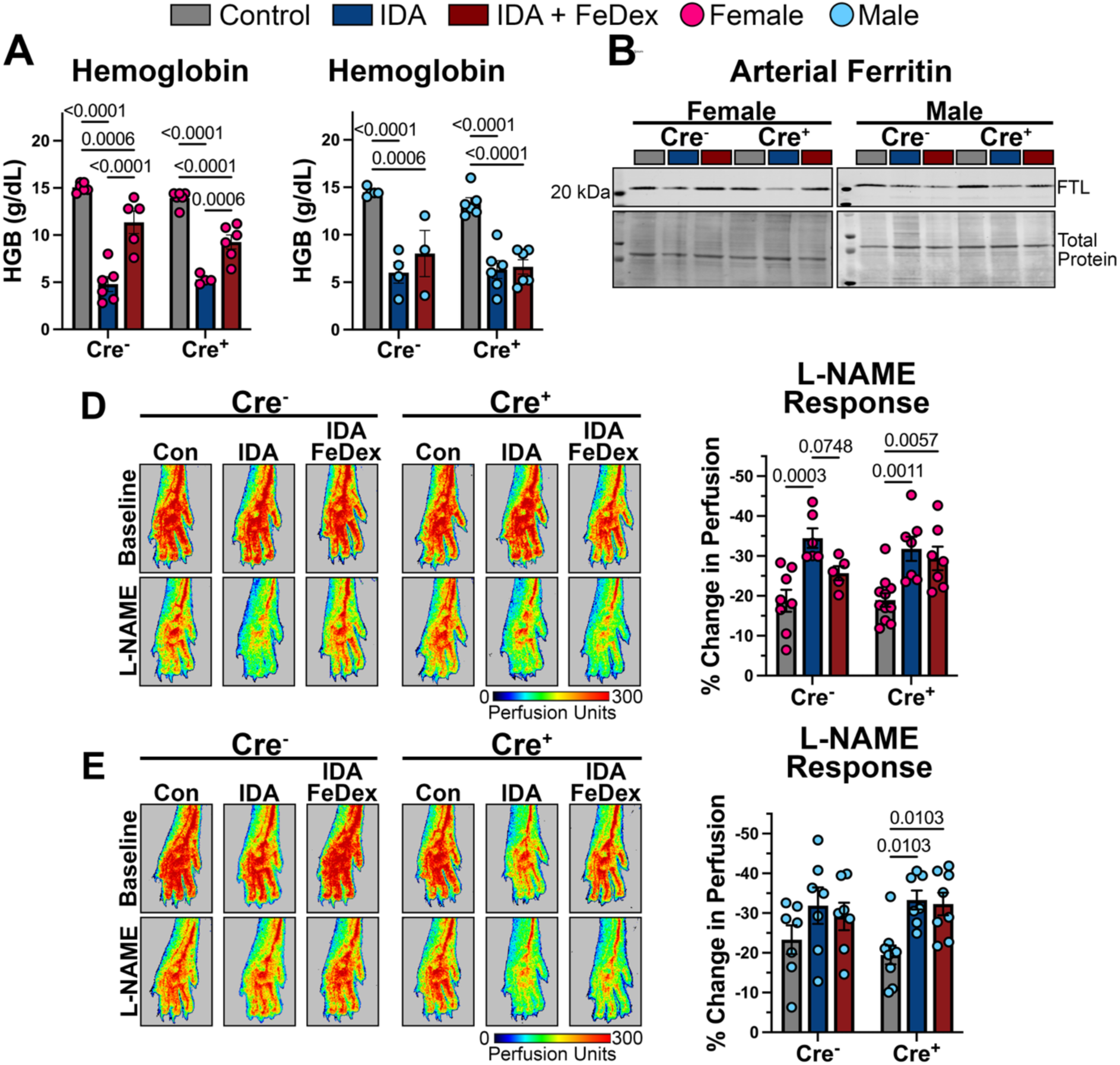
Endothelial iron controls nitric oxide via α-globin. Cdh5-CreER^T2+/-^ Hba1^fl/fl^ mice were fed control (48 ppm Fe) or iron deficient diet (2-6 ppm Fe) beginning at weaning. Tamoxifen was administered intraperitonially at 6 wks for 10 consecutive days. Phlebotomies of 10% blood volume were performed in iron deficient mice at 9 weeks and 11 weeks to aid in the progression of anemia then received a single injection of iron dextran (FeDex; 20 mg/kg) or vehicle (saline) at 11 weeks. All experiments were performed at 13 weeks. (A-B) Blood hemoglobin was measured across all groups. (C) Ferritin light chain (FTL) assessed by western blot in isolated mesenteric arteries. (D-E) Changes in blood flow in hind paws of female (D) and male (E) mice were assessed by laser speckle contrast imaging in response to intraperitoneal injection of the nitric oxide synthase inhibitor, L-NAME. Statistical significance was determined by two-way ANOVA and Holms-Sidac post hoc analysis.

## Discussion

It is well recognized that IDA is associated with perturbed hemodynamics, but the literature investigating these changes has largely neglected direct actions of iron deficiency on the vasculature. Here we demonstrate iron levels vary across the vasculature and directly impact endothelial phenotypes. Furthermore, we identify a role for endothelial Hbα in mediating the increased NO observed in anemia.

The endothelium is specialized into distinct phenotypes that vary along the vascular tree. We used single cell transcriptomic datasets of ECs from mice and humans to investigate if iron varies between endothelial subtypes. We found iron levels to be lowest in arterial ECs and highest in lymphatic ECs. Notably, this pattern did not match the expression of iron dependent proteins across the endothelium suggesting differential iron regulation is not due to increased iron utilization. We therefore investigated the impact of iron chelation on EC phenotypes. We found iron chelation upregulated the expression of arterial markers in human arterial and lymphatic ECs and decreased lymphatic markers in human lymphatic ECs. It is worth noting, not all markers of arterial and lymphatic ECs changed in the hypothesized direction suggesting iron is not sufficient to fully shift EC identity. Iron is likely one of many signals which regulate EC identity. It remains to be studied how iron interacts with or mediates the known effects of VEGF, shear stress, Notch signaling, and cell cycle state on endothelial differentiation^21^.

Arteries themselves are heterogeneous. Large conduit arteries such as the aorta are exposed to high shear stress and have a thick inner elastic lamina to dampen the pulsatile flow generated by the heart. Small resistance arteries are the main source of vascular resistance. These arteries are exposed to lower shear stress and have a thin, discontinuous inner elastic lamina which allows the formation of MEJs by which ECs regulate vascular tone. We modeled both shear stress and MEJs *in vitro* to examine whether they impact endothelial iron. While the presence of MEJs does not change endothelial iron stores or the labile iron pool, exposing cells to shear stress increases intracellular iron consistent with the phenotype observed in conduit artery ECs. These results explain the difference in iron between arterial EC subtypes, but do not explain differences we observed among other vascular segments as iron is highest in the lymphatic endothelium even though shear stress is lowest in these vessels. These differences may be due to other stimuli such as oxygen tension or may be endogenously programed in the cell’s identity. Ultimately, our data suggest the resistant artery endothelium is iron deplete relative to other EC subtypes. Having limited iron stores may make these ECs especially susceptible to iron deficiency. We therefore investigated how the resistance artery endothelium is impacted in IDA.

Increased arterial NO signaling has been previously observed in patients with anemia and attributed to a decreased scavenging of NO by erythrocytes^5^. Free hemoglobin is a well-recognized and potent NO scavenger with a reaction constant of >6×10^7^ M^-1^s^-1^ ^6^. However, *ex vivo* artery preparations demonstrate limited impact of erythrocytes on arterial NO signaling. For example, studies in the rat renal microvasculature have shown minimal scavenging of arterial NO by erythrocytes with the maximum effect at 1% hematocrit^22^. This is well below what is observed in anemia or any physiologic setting. Other studies in pig coronary microvessels found no effect of RBCs (60% hematocrit) on flow mediated dilation, an NO-dependent process^23^. This suggests changes in hematocrit may be insufficient to increase NO signaling. For this reason, we developed a mouse model of IDA in which we can replete vascular iron without fully rescuing the anemia. We did observe a small increase in hematocrit in female mice treated with FeDex, but this is still meaningfully below control mice. To assessed NO signaling, we measured blood flow in the hind paws before and after administration of L-NAME. This method is analogous to the studies performed in the forearm of anemic patients^5^. L-NAME causes vasoconstriction by inhibiting NO synthesis therefore the magnitude of the response to L-NAME can used as a measure of NO signaling. We observed elevated NO signaling in female but not male IDA mice which was rescued by repletion of vascular iron with FeDex. We found IDA did not impact nitrite, nitrate, eNOS expression, or eNOS activation by phosphorylation at S1177 suggesting that NO synthesis was not responsible for the increased NO signaling.

Our lab has previously shown Hbα is expressed in the resistance artery endothelium and inhibits NO signaling by directly binding to eNOS and scavenging NO^7,9,10,24^. We found IDA decreases Hbα in both male and female mice and FeDex rescues Hbα expression in female mice. We further measured the heme-bound portion of Hbα our model and found holo-Hbα to be at the detection limit of the assay for females but easily detectable in males. IDA decreased holo-Hbα in males which was not rescued by repleting vascular iron whereas holo-Hbα was increased in iron replete females. Based on these data, we hypothesized changes in NO signaling may be due to the regulation of holo-Hbα. To test this hypothesis, we utilized endothelial-specific *Hba1* knockout mice and found deletion of endothelial Hbα prevented the regulation of NO signaling by FeDex in female mice demonstrating iron regulates NO signaling through endothelial Hbα. In male Cre^+^ mice, IDA increased NO which was not rescued by FeDex phenocopying the female Cre^+^ mice. Based on these findings, we conclude loss of Hbα is necessary for increased NO signaling in IDA in both males and females. However, the increased NO signaling does not occur in males because they have a larger pool of holo-Hbα which is not fully lost in IDA.

Anemia has long been recognized to regulate hemodynamics by decreasing systemic vascular resistance^5^, but the role of endothelial iron deficiency in these changes has been overlooked. Our data demonstrate resistance arterial iron and its regulation of Hbα play a critical role in modulating hemodynamics in IDA. Furthermore, the work demonstrates a novel role for Hbα mediating sex differences in vascular biology. Although there were similar levels of iron deficiency in males and females, female mice were more sensitive to vascular iron deficiency due to their nearly absent holo-Hbα. In humans, the impact of this sensitivity may be potentiated by the high prevalence of IDA in females relative to males^25^. Understanding the role of iron in vascular biology in males and females will help elucidate the hemodynamic consequences of one of the most common nutrient deficiencies^26^.

## Acknowledgements

NIH HL007284 (L.S.D., S.A.L, W.S.J., and M.A.L); NIH HL172605 and AHA 25CDA1444441 (L.S.D); HL176103 (S.A.L.); AHA 25PRE1375860 (W.S.J.); AHA 25PRE1376958 (Z.J.); AHA915176 (M.A.L); NIH HL137112 and HL171997 (B.E.I.); UVA Comprehensive Cancer Center Farrow Fellowship (B.O.)

