## Supplemental Figures for "Arterial iron regulates vasodilation during anemia via endothelial holo-alpha globin"

Supplemental figures: 4

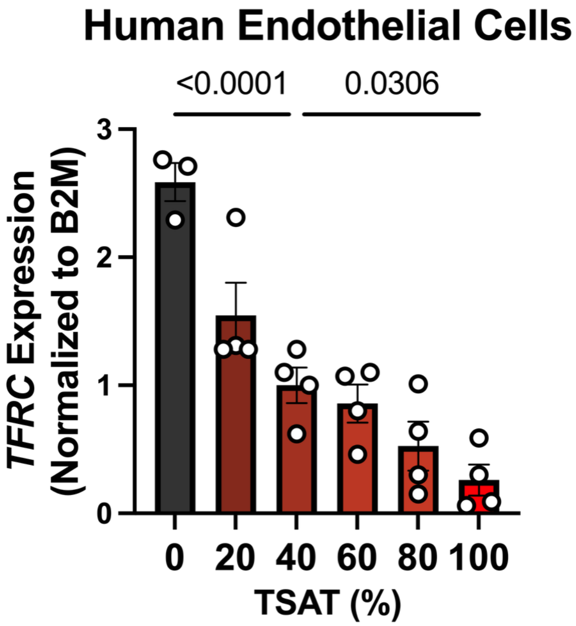


**Supplemental Figure 1. *TFRC* mRNA inversely correlates with endothelial iron.** *TFRC* was assessed by qPCR in endothelial cells treated with transferrin at varying degrees of iron saturation. Statistical significance was determined by one-way ANOVA with Holms-Sidac post hoc test.

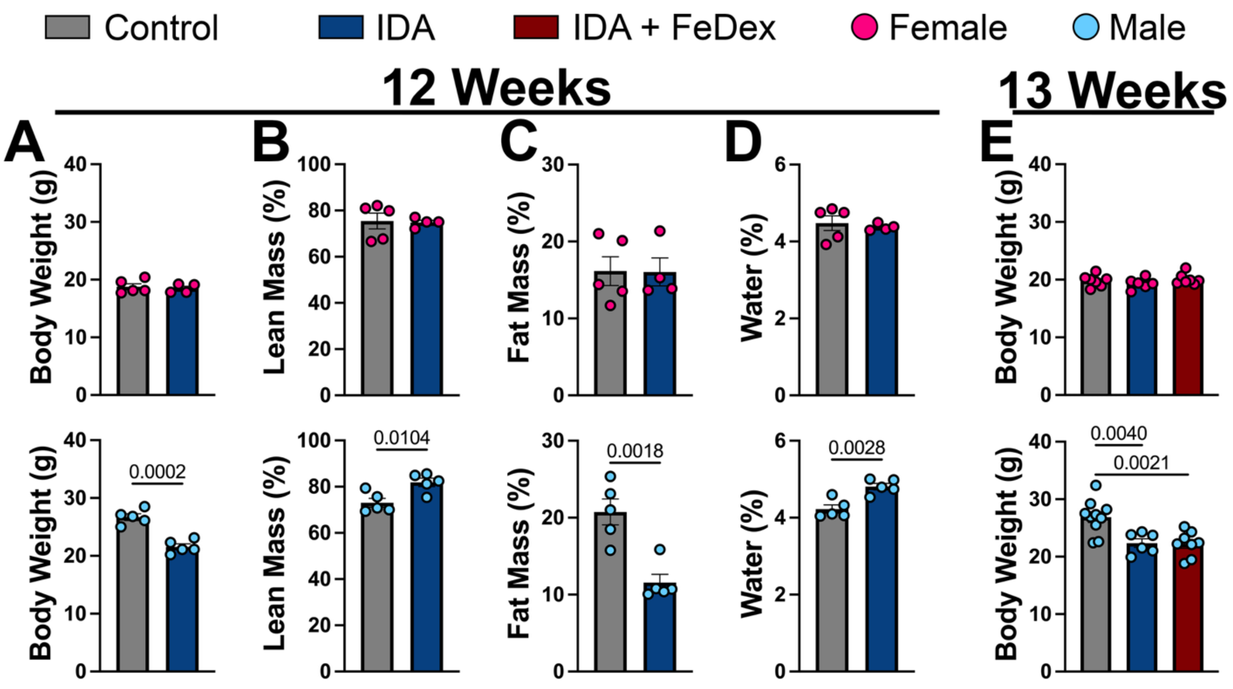


**Supplemental Figure 2. Body weight and mass distribution in mice with iron deficiency anemia.** Body mass was assessed at 12 and 13 weeks. Body mass distribution was assessed by echoMRI at 12 weeks. Statistical significance was determined by Student's t-test or one-way ANOVA with Holms-Sidac post hoc test.

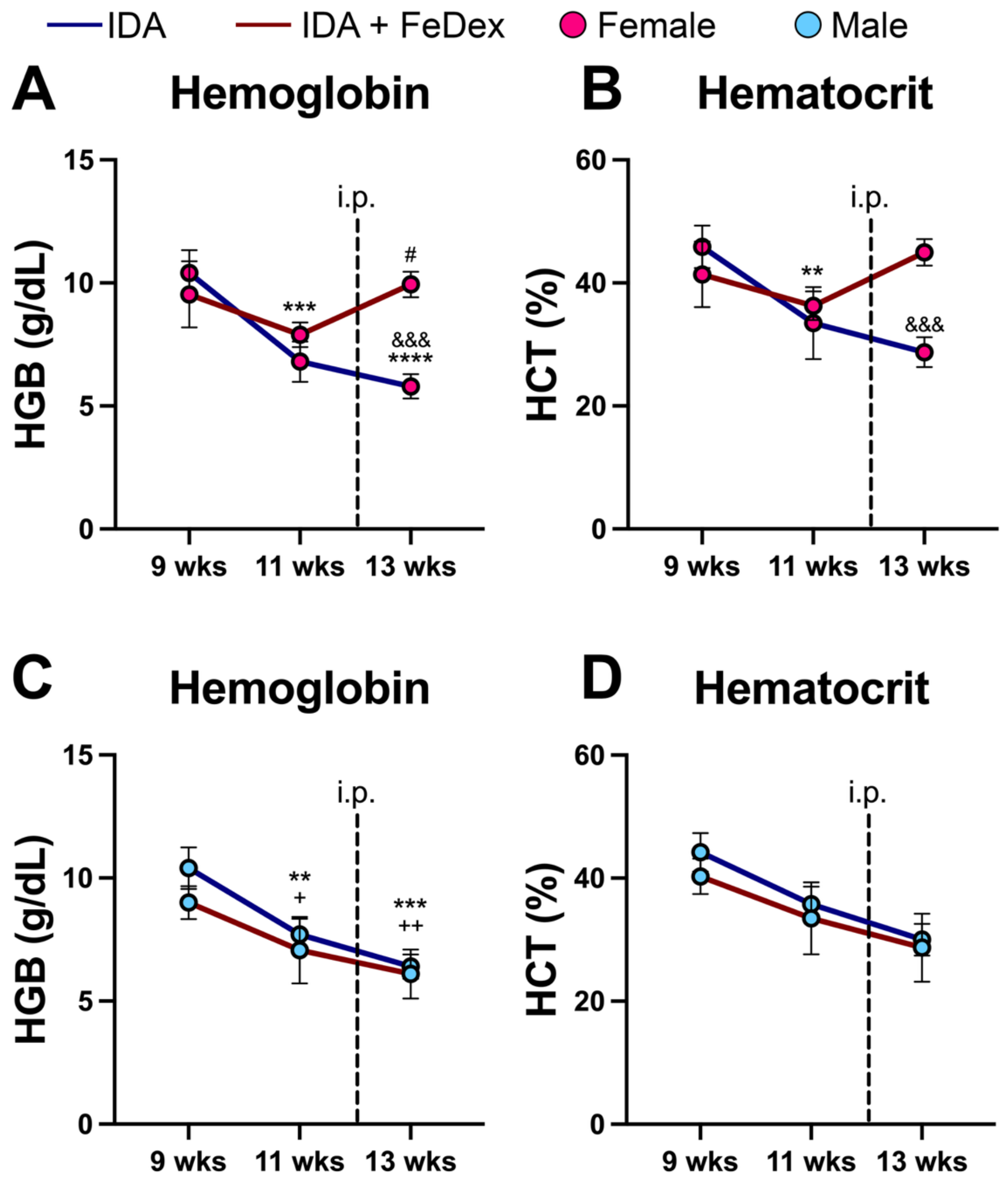


**Supplemental Figure 4. Progression of anemia during the iron deficiency anemia protocol.** Hemoglobin and hematocrits were assessed at 9, 11, and 13 weeks in male and female mice. Statistical significance was determined by two-way ANOVA with Holms-Sidac post hoc test.

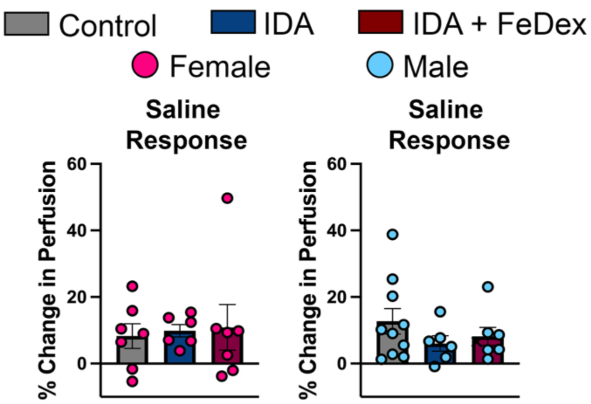


**Supplemental Figure 5. Laser speckle contrast imaging vehicle control.** Changes in blood flow in response to saline were assessed in the hind paws of mice. Statistical significance was determined by one-way ANOVA with Holms-Sidac post hoc test.
